# Protocol for in vivo DNA-RNA hybrid immunoprecipitation sequencing and analysis from frozen mammalian tissues

**DOI:** 10.64898/2026.04.06.716701

**Authors:** Hassan Massalha, Cameron J. Chee, Julia S. P. Mawer, Francesco Puzzo, Magdalena P. Crossley

## Abstract

DNA-RNA hybrids (R-loops) form transiently on the genome and regulate cellular homeostasis. They also influence genome editing outcomes, highlighting their therapeutic potential in vivo. This protocol enables high-resolution mapping of DNA-RNA hybrids directly from frozen mouse tissues. Following tissue homogenisation and lysis, genomic DNA is extracted, digested and DNA-RNA hybrids are isolated using the hybrid-specific S9.6 monoclonal antibody. The purified hybrids are then processed for whole-genome sequencing to generate R-loop profiles. For complete details on the use and execution of this protocol, please refer to Puzzo, Crossley et al^1^.

**graphical abstract.**
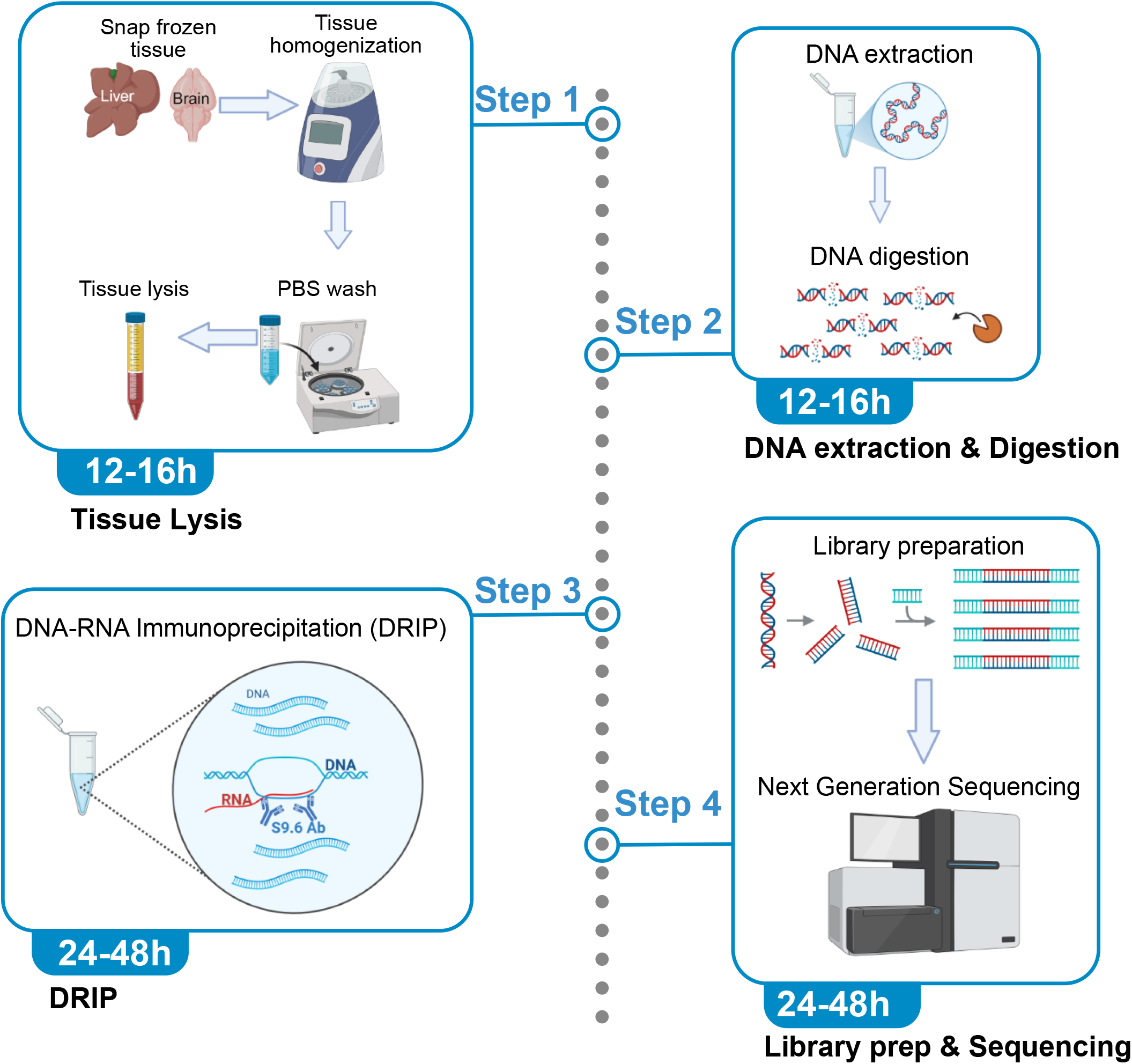

## Innovation

This protocol describes the first implementation of DRIP-seq directly on intact mammalian tissue. The previous DRIP-seq studies performed on tissue samples have generally relied on freshly isolated material followed by genomic DNA extraction from dissociated single cells. Our approach described herein enables the direct processing of snap-frozen tissue, allowing samples to be conveniently collected, stored, and analyzed without the requirement for immediate processing^1^.

By enabling genome-wide mapping of RNA-DNA hybrids (R-loops) directly from frozen tissues, this method expands R⍰loop profiling into physiologically relevant in vivo contexts beyond cultured cell lines, providing a practical framework for studying the distribution and potential functional roles of R⍰loops in animal models of disease. In addition, we provide bioinformatic workflows to directly compare matched tissue and cell⍰line DRIP⍰seq datasets, broadening the applicability of DRIP⍰seq to tissuel⍰based studies and enabling systematic investigation of R⍰loop dynamics across diverse biological and pathological settings.

### Institutional permissions

Animal experiments described in the present protocol were conducted and approved by the Administrative Panel on Laboratory Animal Care of Stanford University relative to the publication Puzzo F., Crossley M.P., et al.^1^

### Pre-experimental preparation

The following procedure has been validated for snap-frozen organs (liver and brain) collected upon transcardiac perfusion.

1. Deeply anesthetize the mice.
2. Transcardially perfuse animals with room-temperature Dulbecco’s phosphate-buffered saline (DPBS)^2,3^.
3. Quickly rinse tissues in 10 mL DPBS inside a 10 cm^2^ Petri dish, transfer to 2 mL tubes, and immediately snap-freeze on dry ice.
4. Store samples at -80°C until further use.

The following phase separation gel protocol for making phase lock tubes was validated for use in DNA extraction, digestion and DNA-RNA hybrid immunoprecipitation.

**Note:** Commercially available phase lock tubes have been discontinued, so homemade phase lock tubes have been provided as an alternative.

1. Add High Vacuum Grease and 10% Silicon Dioxide (SiO_2_) to a high quality ziploc bag, then mix thoroughly using a combination of manual kneading and rolling **(Figure 1A, 1B)**.
2. Cut a corner of the plastic bag, dispense the phase separation gel into syringes and autoclave **(Figure 1C)**.
3. After cooling to room temperature, aliquot 1.5 mL of the gel into 15 mL tubes (100 μl into 2 mL tubes), spin down at 3000x g for 1 min (16,000x g for 2 mL tubes), and store upright at room temperature until needed **(Figure 1D)**.

**Figure 1:**
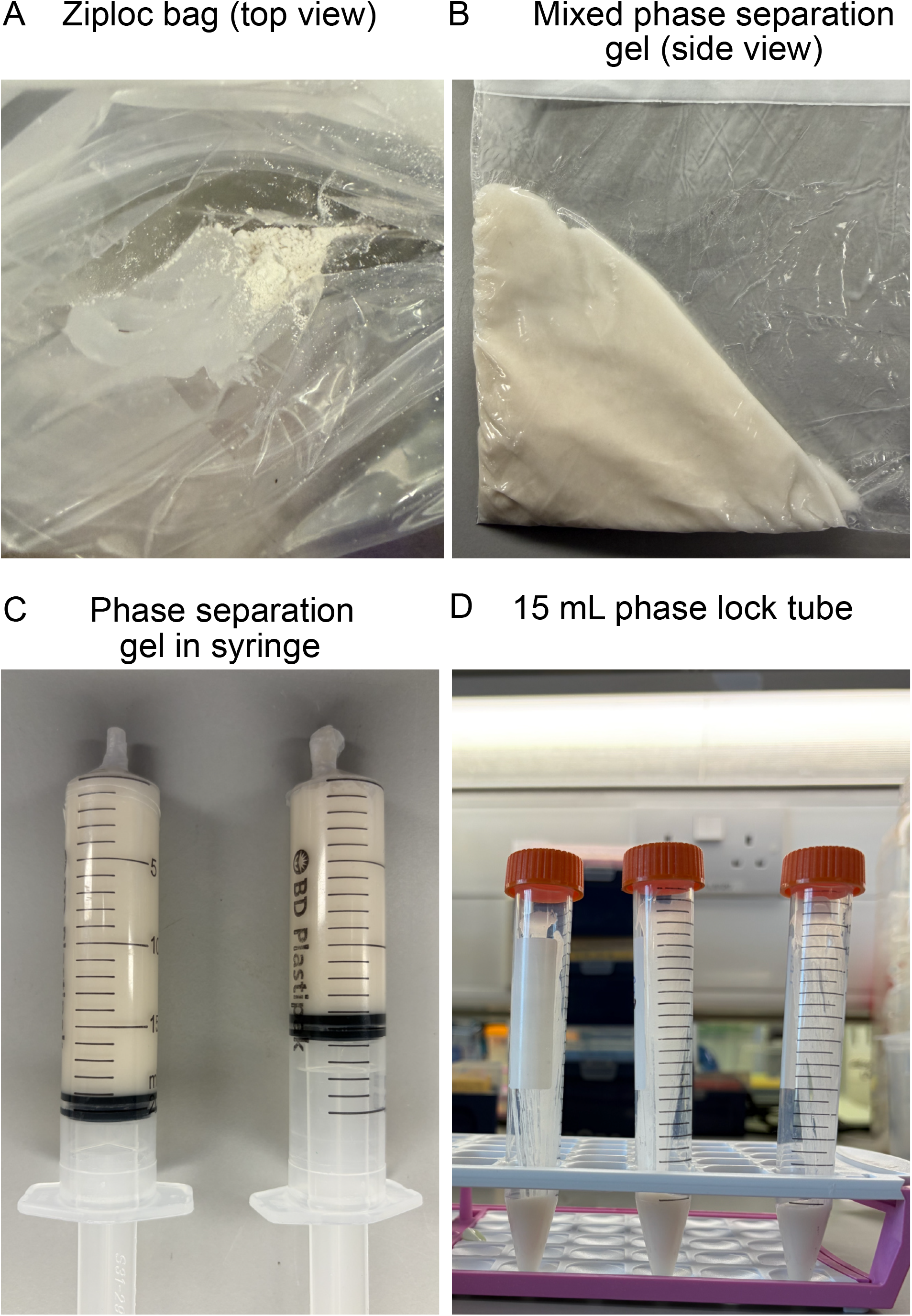
Preparation of phase lock tubes. (A) 10% SiO_2_ vacuum grease before mixing. (B) 10% SiO_2_ vacuum grease mixed into a smooth paste. (C) Phase separation gel in syringes. (D) 15 mL phase lock tubes stored upright at room temperature.

**Table 1:**
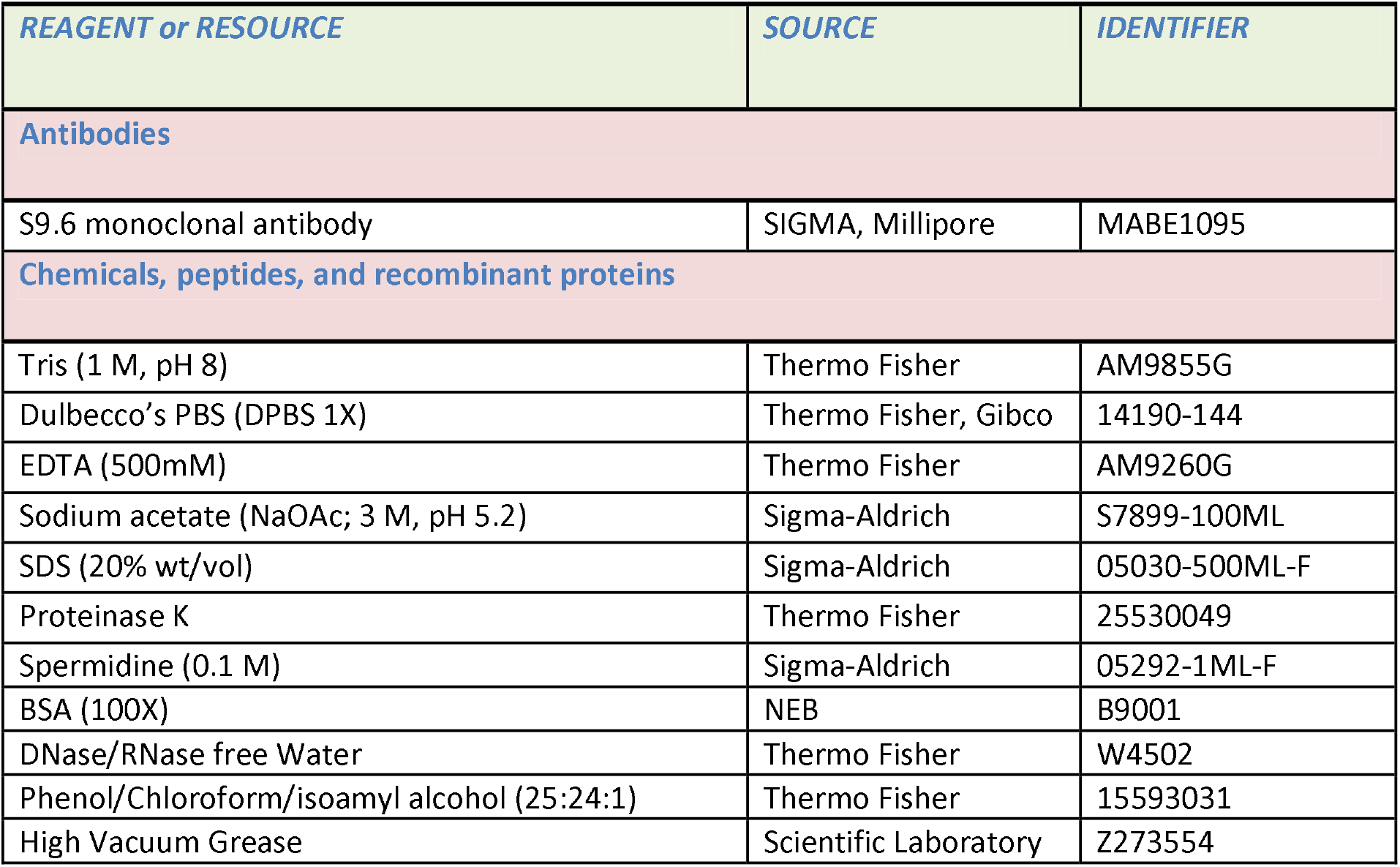
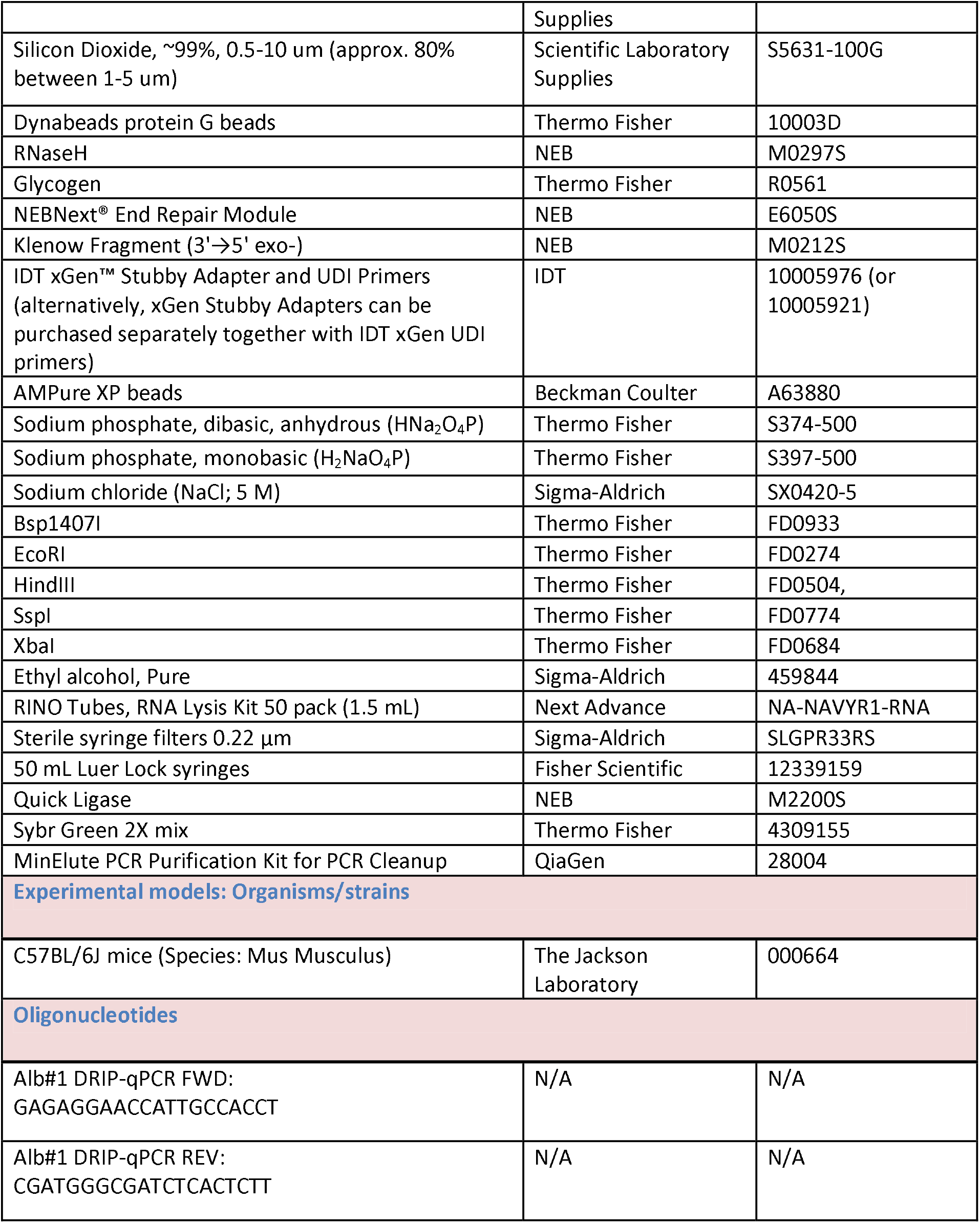
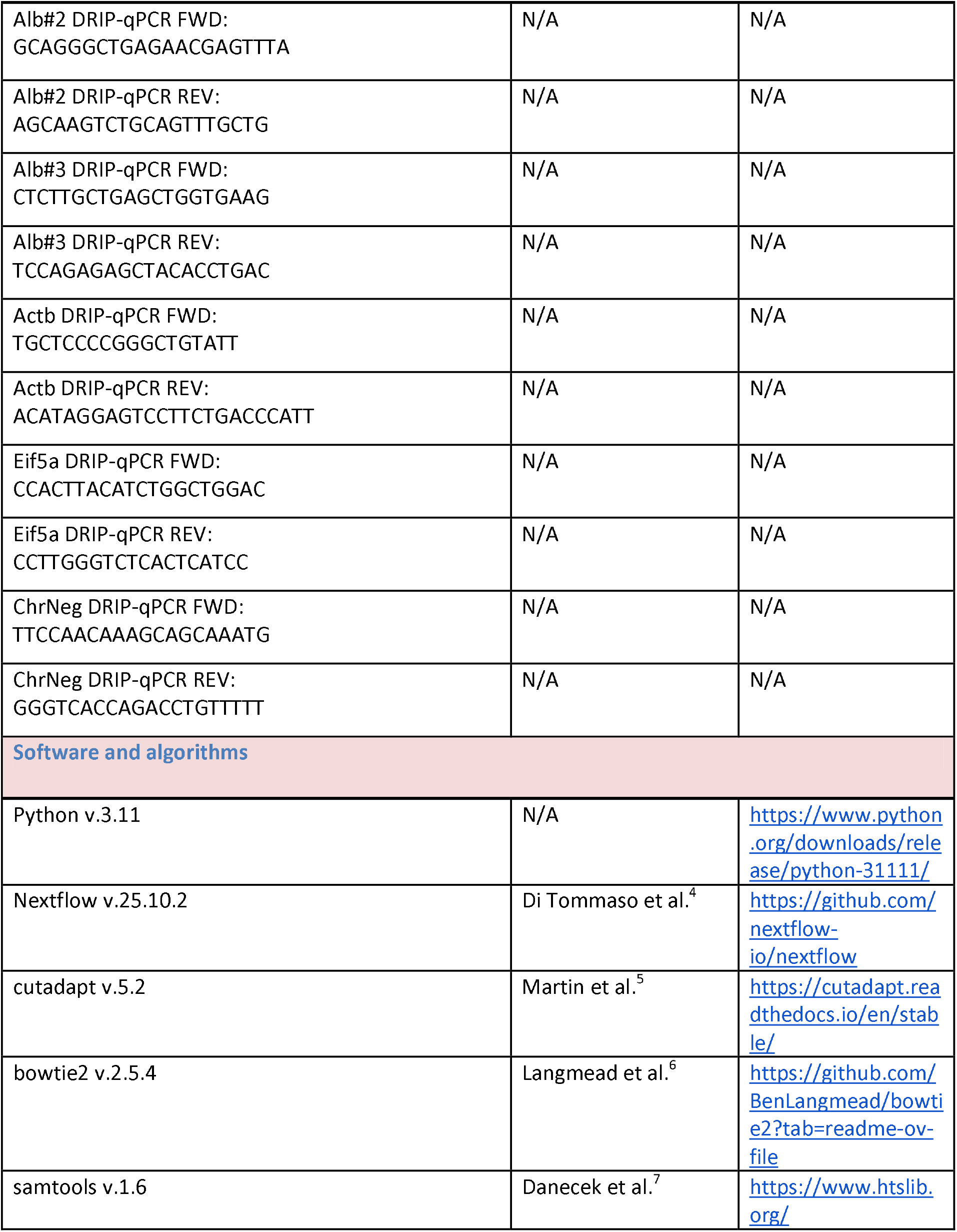
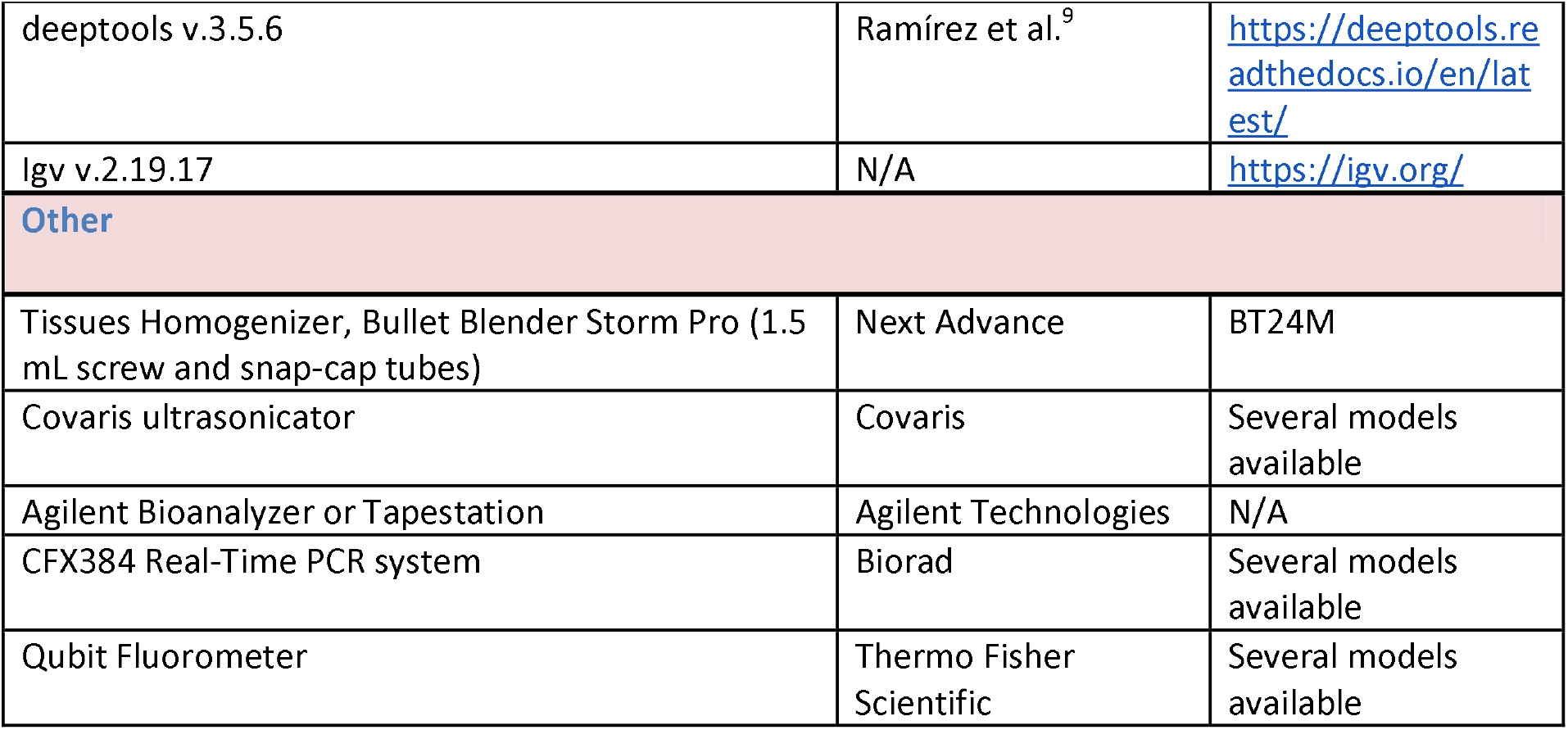
Key resources table.

## Materials and equipment

### Phase separating gel

Materials and equipment Phase separating gel
| Reagent | Final concentration | Quantity or Volume |
| --- | --- | --- |
| High Vacuum Grease | N/A | 5 g |
| Silicon Dioxide (~99% 0.5-10 $\mu\text{m}$) | 10% w/w | 0.5 g |
| Total |  | 5.5 g |

### TE buffer

| Reagent | Final concentration | Quantity or Volume |
| --- | --- | --- |
| Tris (1 M, pH 8) | 10 mM | 500 $\mu\text{L}$ |
| EDTA (500 mM) | 1 mM | 100 $\mu\text{L}$ |
| DNase/RNase-free water |  | 49.4 mL |
| Total | n/a | 50 mL |

### Sodium phosphate 1 M, pH 7

| Reagent | Final concentration | Quantity or Volume |
| --- | --- | --- |
| Sodium phosphate, monobasic | 2 M | 39 mL |
| Sodium phosphate, dibasic, anhydrous | 2 M | 61 mL |
| DNase/RNase-free water |  | 100 mL |
| Total | 1 M | 200 mL |

**Note:** If this solution crystallizes or precipitates, warm it up to help resolubilize it. The 1 M sodium phosphate, pH 7, solution can be stored indefinitely at room temperature.

### 10X DRIP binding buffer

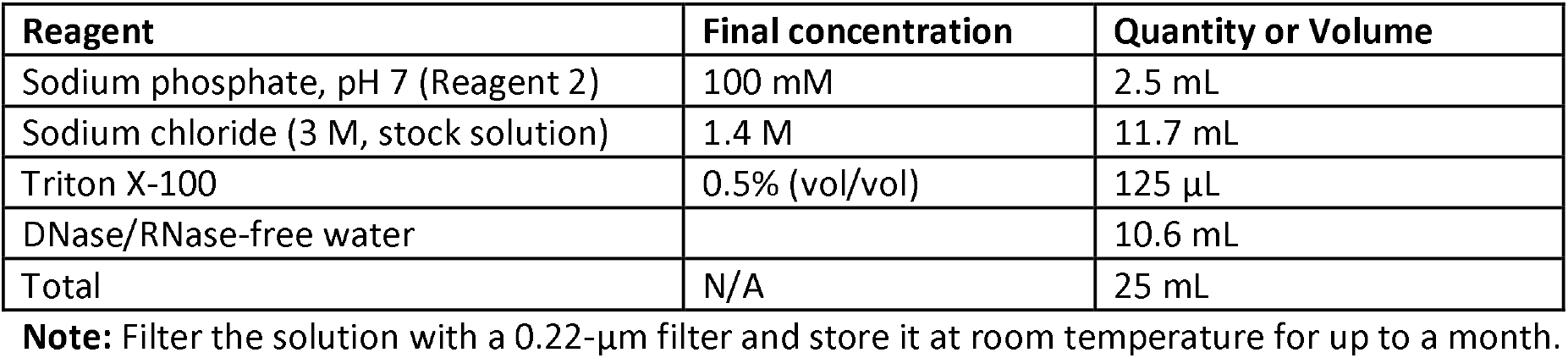

| Reagent | Final concentration | Quantity or Volume |
| --- | --- | --- |
| Sodium phosphate, pH 7 (Reagent 2) | 100 mM | 2.5 mL |
| Sodium chloride (3 M, stock solution) | 1.4 M | 11.7 mL |
| Triton X-100 | 0.5% (vol/vol) | 125 $\mu\text{L}$ |
| DNase/RNase-free water |  | 10.6 mL |
| Total | N/A | 25 mL |

**Note:** Filter the solution with a 0.22- $\mu\text{m}$ filter and store it at room temperature for up to a month.

### DRIP elution buffer

| Reagent | Final concentration | Quantity or Volume |
| --- | --- | --- |
| Tris (1 M, pH 8) | 50 mM | 1.25 mL |
| EDTA (500 mM) | 10 mM | 500 $\mu\text{L}$ |
| SDS (20% w/vol) | 0.5% (vol/vol) | 625 $\mu\text{L}$ |
| DNase/RNase-free water |  | 22.6 mL |
| Total | N/A | 25 mL |

**Note:** This solution can be stored for up to a year at room temperature.

## Step-by-step method details

All centrifugation steps are performed at room temperature, unless otherwise specified.

### Tissue lysis

#### Timing: 12-16h

This procedure describes how to perform the lysis of the snap-frozen organ for the next step of DNA extraction.

1. Retrieve snap-frozen liver tissue (the mice must be cardiacally perfused with PBS) from -80°C and allow it to thaw on ice for 30 min. **Note:** The cardiac perfusion of the animal will avoid and minimize contamination from blood cells.
  a. Transfer tissue into RINO tubes (2-3 tubes per liver). **Note:** If stored as a single frozen piece, section it on dry ice into fragments small enough to fit the tubes.
  b. Add 500 µL ice-cold sterile DPBS to each tube and homogenize using a Next Advance homogenizer for 5 min at speed 8.
  c. Pool all homogenates into a single 2 mL tube to ensure representation of the entire organ. **Note:** The homogenization of the whole liver is important to avoid zonal metabolic and transcriptional bias due to the partial organ homogenization.
2. Dilute 250 µL homogenate in 5 mL PBS and centrifuge at 300x g for 5 min at 4°C . Repeat the wash twice. **Note:** The rest of the homogenate can be aliquoted and stored at -80°C for up to one year. It can be used for either other DRIP studies or different experiments (the sample is resuspended in DPBS and therefore can be utilized for nucleic acids or protein extraction).
3. After the last wash, resuspend the final pellet containing the hepatic cells in 1.6 mL of TE Buffer.
4. Add 8 µL Proteinase K and 50 µL of 20% SDS. **Note:** Gently mix by inverting the tube 3-4 times.
5. Incubate the tube at 37°C overnight in a water bath or an incubator.

### DNA extraction and digestion

#### Timing: 12-16h

This section describes the extraction of the genomic DNA from tissues gently lysed overnight. It also explains how to fragment the DNA by using a specific cocktail of restriction enzymes.

6. Centrifuge 15 mL homemade phase lock tubes at 3,000 xg for 5 min.
  a. Transfer the lysate incubated overnight at 37°C into the 15 mL phase lock tube
  b. Add an equal volume (1.6 mL) of phenol:chloroform:isoamyl alcohol (25:24:1).
  c. Invert 4-5 times and centrifuge at 1,500 xg for 5 min.
  **Note:** After this step, the supernatant should be clear. If not, spin for an additional 5 min.
7. Add a 1/10 volume of 3 M NaOAc, pH 5.2 (160 µL), and 2.5 volumes of 100% (vol/vol) ethanol (4 mL) to a new 15 mL tube.
  a. Pour in the DNA (the top aqueous phase) from the phase lock gel tube and mix the DNA and ethanol gently by inverting the tube until the DNA fully precipitates (visible by eye as a white precipitate).
  b. Wait for 10 min until all DNA is precipitated.
  c. Spool the DNA out of the mixture with a 1000 µL cut tip and transfer it to a clean 2 mL tube.
  **Note:** Do not pellet DNA by centrifugation to minimize RNA contamination and preserve RNA-DNA hybrids.
8. Wash the DNA by adding 1 mL of 80% (vol/vol) ethanol freshly prepared, gently inverting the tube 2-3 times and letting it stand for 3 min.
  a. Carefully discard the supernatant; avoid pipetting the DNA.
  b. Repeat this step twice.
  **Note:** Very carefully remove as much ethanol as possible by pipetting after the last wash.
9. Allow to air dry completely while inverting tubes (to give a clear pellet). **Note:** This usually takes ∼10 min. If more cells are lysed this can take longer. Marking the location of the DNA on the tube can help to locate it once it has dried.
  a. Add 125 μL of TE buffer directly to the DNA pellet and carefully dislodge the cell pellet from the side of the tube with a pipette tip.
  b. Keep on ice for 1 h.
  c. Gently resuspend the DNA by pipetting two or three times with a 200 μL cut tip.
  d. Leave on ice for at least another hour before starting the restriction enzyme digest.
  **Note:** Avoid vortexing and overpipetting; maintaining the DNA viscosity will better preserve intact RNA-DNA hybrids.
10. Digest whole solution ∼125 μL of extracted genomic DNA using a cocktail of restriction enzymes: BsrGI (Bsp1407I), EcoRI, HindIII, SspI and XbaI (5 μL/enzyme in each reaction). Incubate overnight at 37°C.
  a. For 200 μL final reaction volume: 125 μL DNA + 5 μL x 5 enzymes (total 25 μL) + 20 μL enzyme Buffer 10X + 27 μL DNase/RNase-free water + 1.5 μL BSA + 1.5 μL Spermidine.
  **Note:** Spermidine should be added last at the proper concentration; excess spermidine can cause DNA precipitation. After overnight incubation, the viscosity of the mixture should have disappeared if the digestion is complete.
11. Spin a 2 mL phase lock tube for 1 min at 16,000 xg to pellet the gel.
  a. Gently pipette the DNA from the previous step into the phase lock gel tube.
  b. Add 100 μL of DNase/RNase-free water and 1 volume (300 μL) of phenol/chloroform/isoamyl alcohol (25:24:1).
  c. Gently invert the tube 4 to 6 times and spin down at 16,000 xg for 10 min.
12. In a clean 1.5 mL tube, mix 1.5 μL of glycogen, 1/10 volume of 3 M NaOAc, pH 5.2 (30 μL), and 2.5 volumes of 100% (vol/vol) ethanol (750 μL).
  a. Gently pipette in the DNA (top aqueous phase) from the phase lock gel tube and mix by inverting 4 to 6 times.
  b. Incubate for at least 2 h at -20°C (overnight incubation will enhance the DNA recovery) to increase precipitation yields.
13. Spin at 16,000 xg for 35 min at 4°C.
  a. Discard the supernatant.
  b. Add 200 μL of freshly made room-temperature 80% (vol/vol) ethanol.
  c. Spin for 10 min at 16,000 xg at 4°C and discard the supernatant.
14. Air dry the pellets for 10 min at room temperature until they become transparent, depending on the DNA concentration, and resuspend in 50 μL of TE buffer.
  a. Leave the tube on ice for 30-60 min and then gently resuspend.
  b. Measure the concentration (OD260) of your DNA on a spectrophotometer. Typically, you should expect a concentration of at least ∼1 μg/μL.
  **Note:** To resuspend the DNA, do not vortex the tube or overpipette. Leave on ice longer or add an additional 50 μL TE buffer to help resuspension. **Note:** A specificity control is provided by including a sample that is pre-treated with RNaseH to remove RNA-DNA hybrids from the mixture. Store the DNA that is not treated with RNaseH at 4°C overnight before starting the IP the day after. (It can be stored for a few days at 4°C). Alternatively, the DNA can be stored at -80°C for up to a month.
15. Dilute 4-10 μg of digested DNA with 5-10 μL RNaseH in 1X RNase H digestion buffer. **Note:** The final reaction volume should be at least 10 times the volume of RNaseH added to dilute the enzyme storage buffer (i.e., 8 μg of DNA + 1X RNaseH buffer + 8 μL RNaseH + up to 80 μL of RNase/DNase-free water).
  a. Incubate overnight at 37°C.

Proceed to S9.6 immunoprecipitation step below.

### DNA-RNA immunoprecipitation (DRIP)

#### Timing: 24-48h

This section describes how to prepare the sample and perform the immunoprecipitation of the DNA-RNA hybrids (R-loops) using the S9.6 monoclonal antibody.

16. Samples treated overnight with RNaseH.
  a. Add 20 μL of RNase/DNase-free water to the RNaseH solution up to a final volume of 100 μL.
  b. Add the sample to a 2 mL phase lock tube with 1 volume (100 μL) of Phenol:Chloroform:Isoamyl alcohol (25:24:1) and invert the tubes 4-5 times.
  c. Spin the phase lock tubes for 10 min at 16,000 xg.
  d. Add the top aqueous phase to a clean tube containing 1.5 μL of glycogen, 1/10 volume of 3 M NaOAc, pH 5.2 (10 μL), and 2.5 volumes of 100% (vol/vol) ethanol (250 μL) and invert the tubes 4-5 times.
  e. Incubate for at least 2 h at -20°C or at least 1h at -80°C to precipitate the DNA. Longer incubation times can be used, if practical.
  f. Spin at 16,000 xg for 35 min at 4°C.
  g. Discard the supernatant and add 200 μL of room-temperature 80% (vol/vol) ethanol.
  h. Spin for 10 min at 16,000 xg at 4°C and discard the supernatant.
  i. Air dry the pellets for 10 min at room temperature until they become transparent.
  j. Add 50 μl TE and leave the tubes on ice for 30 min to resuspend the pellets.
17. Dilute 8 μg DNA in 500 μL TE.
  a. Save a 1/10 volume for each tube (50 μL) to use as input for qPCR. Store at -20°C.
18. Add 50 μL of 10X DRIP binding buffer and 20 μL of the S9.6 antibody to the 450 μL of diluted DNA.
  a. Incubate the tubes for 14-17 h at 4°C while gently inverting on a mini-tube rotator.
19. For each tube, wash 50 μL of the Dynabeads (equilibrate beads at room temperature for 10 min) with 700 μL of 1X DRIP binding buffer (1X DRIP binding buffer is made by diluting the 10X DRIP binding buffer in TE buffer) by inverting the tubes on a mini-tube rotator for 10 min at room temperature.
  a. Put the tubes onto a magnetic rack for 1 min and discard the supernatant.
  b. Repeat this step once for a total of 3 washes.
20. Add DNA:antibody complexes to prewashed beads, incubate 2 h at 4°C on a rotating shaker.
  a. Spin down beads briefly to remove liquid from the lid and sides of the tube.
  b. Apply to the magnet and discard the supernatant.
21. Wash the beads:antibody complexes with 700 μl of 1X binding buffer,
  a. Invert 5-10 min at room temperature on a rotating shaker.
  b. Spin down and discard supernatant as previously described.
  c. Repeat twice for a total of 3 washes.
22. Add 250 μL of DRIP elution buffer and 8 μL of proteinase K (20 mg/mL) to the beads.
  a. Seal the tubes with Parafilm to avoid any leaking.
  b. Incubate with rotation at 55°C for 45 min in a temperature-controlled rotating oven.
23. Put the tubes onto a magnetic rack for 1 min.
  a. Meanwhile, spin 2 mL phase lock tubes for 1 min at 16,000x g to pellet the gel.
  b. Transfer the supernatant of each tube to the 2 mL phase lock tube.
  c. Add 1 volume (250 μL) of phenol/chloroform/isoamyl alcohol (25:24:1).
  d. Invert gently 4-5 times.
  e. Spin down at 16,000x g for 10 min at room temperature.
24. In a clean 1.5 mL tube, mix 1.5 μL of glycogen, 1/10 volume of 3 M NaOAc, pH 5.2 (30 μL), and 2.5 volumes of 100% (vol/vol) ethanol (750 μL).
  a. Gently pipette in the DNA (top aqueous phase) from the phase lock tube.
  b. Mix by inverting four to six times.
  c. Incubate for at least 2 h (overnight incubation will increase the DNA recovery) at - 20°C to increase precipitation yields.
25. Spin at 16,000x g for 35 min at 4°C.
  a. Discard the supernatant.
  b. Add 200 μL of room temperature 80% (vol/vol) ethanol.
  c. Spin for 10 min at 16,000x g at 4°C and discard the supernatant.
26. Air-dry the pellets for 10-15 min.
  a. Add 50 μL of TE buffer to each tube.
  b. Leave the tubes on ice for 15-30 min and gently resuspend.
  **Pause point:** The immunoprecipitated DNA can be stored at –80°C for up to six months.
27. Check the DRIP efficiency by qPCR.
  a. Dilute samples 1:5 in RNase/DNase-free water and use 3 µL for qPCR

**Note:** Before proceeding, it is advised to make sure that the DRIP procedure worked. To this end, use two negative and two to three positive loci and measure their immunoprecipitation as a fraction of input DNA. The RNaseH-treated sample should lead to very low yields (>90% reduction in DRIP efficiency).

##### DRIP-qPCR analysis

To calculate the percentage of input, apply the following formula for each locus: % input = 100*2^(Ct input(corrected) - Ct DRIPed DNA), where Ct (cycle threshold) input(corrected) = (Ct input - log2(10)) (we subtract log2(10) because the input represents 1/10th of the immunoprecipitated DNA).

### Library preparation

#### Timing: 3-4h

This procedure describes the method to prepare libraries for the DRIP sequencing and their quality control using Bioanalyzer instrumentation.

28. Sonicate the DNA to achieve a peak fragment size of 300 base pairs (bp). **Note:** For the purposes of this protocol, a Covaris S220 with Covaris microtubes (Fisher, 520045) is used with the following sonication parameters: Intensity: 4; C/B: 200; DF: 10%; Time: 30 s
29. End Repair the sheared DNA.
  a. Add 2.5 μL NEBNext end Repair enzyme + 5 μL NEB 10X buffer + 2 μL ATP to the 40.5 μL DNA for a total volume of 50 μL.
  **Note:** Do it in 500 μL tubes; pipet up and down to mix and incubate at room temperature for 30 min.
30. Clean up the DNA with Minelute QIAGEN Column.
  a. Add 250 μL of buffer PB.
  b. Add 10 μL of 3 M Sodium Acetate.
  c. Elute in 34 μL; wait 1 min before spinning.
31. A-tailing:
  a. Dilute dATP to 1:100 (from 100 mM stock).
  b. Add 5 μL NEB buffer 2 + 10 μL 1 mM dATP + 1 μL NEB Klenow EXO to the 34 μL DNA for a total volume of 50 μL.
  c. Mix and incubate for 30 min at 37°C (in an incubator).
  **Note:** Use PCR tubes for this step.
32. Clean up with Minelute QIAGEN Column.
  a. Add 250 μL of buffer PB.
  b. Add 10 μL of 3 M Sodium Acetate.
  c. Elute in 12 μL, wait 1 min before spin elution.
33. Ligate adapters
  a. To 12 μL A-tailed DNA (∼20-50 ng), add 2.0 μL xGen Stubby Adapter (3.0 μM, 1:5 diluted in RNase/DNase-free water) + 15 μL 2X NEB Quick Ligase Reaction Buffer + 1 μL NEB Quick Ligase for a total volume of 30 μL.
  b. Incubate at room temperature for 15 min.
  **Note:** Use nuclease-free PCR tubes for this step. **Note:** This protocol was originally published using Illumina Prep2Seq indexed adapters. These are now no longer commercially available, and we provide an alternative with Y-type adapters (xGen Stubby Adapters) without indexes, followed by UDI index addition at the PCR step. Library preparation can also be performed using a single kit for the whole process (for example NEBNext Ultra II DNA library kit). The modular library preparation workflow provided here is a more flexible, readily optimizable, and cheaper alternative. **Note:** For lower DNA input amounts (10-20 ng), using 2.0 μL of 1.5 μM adapters may be more appropriate and result in fewer adapter dimers in the libraries (see expected outcomes).
34. Do a two-sided size selection with Ampure beads. **Note:** Do a 1:1 Ratio with Ampure beads (leave it 10 min at room temperature before use).
  a. Add 70 μL of EB buffer to the 30 μL of DNA.
  b. Add 100 μL of Ampure beads, vortex, spin down quickly, and incubate for 15 min at room temperature.
  c. Apply to the magnet for 2 min or until the liquid clears completely, and <u>*discard the supernatant*</u>.
  d. Wash 2 times with 200 μL fresh 80% EtOH (important to be made fresh), and dry for 10 min (beads should start “cracking”).
  e. Add 11 μL EB buffer, vortex and incubate for 5 min at room temperature.
  f. Apply to the magnet, wait for 2 min or until the liquid clears completely, and <u>*take the supernatant*</u>.
  g. Repeat 1:1 Ratio with Ampure beads.
  h. Add 90 μL of EB buffer to 10 μL of DNA (from previous Step).
  i. Add 100 μL of Ampure beads, vortex and spin down quickly.
  j. Incubate for 15 min.
  k. Apply to the magnet, wait for 2 min or until the liquid clears completely, and <u>*discard the supernatant*</u>.
  l. Wash 2 times with fresh 80% EtOH (important to be made fresh), and dry for 10 min (beads should start “cracking”).
  m. Add 11 μL EB, vortex, and incubate for 5 min at room temperature.
  n. Apply to the magnet, wait for 2 min or until the liquid clears completely, and <u>*take the supernatant*</u>.
35. Run PCR. **Note:** Try to use Qubit HS to estimate DNA quantity for the qPCR. Determination of the number of PCR cycles needs to be done by qPCR **(Figure 2)**.
  a. qPCR reaction setup: 2 μL of adapter-ligated DNA (can also take 1 μL DNA dilute 1:10 and do three replicates per sample, then subtract 3 cycles at end of fitpoint) + 0.5 μL xGen UDI Primer Pair + 7.5 μL of Sybr Green Master Mix + 5 μL of RNase/DNase-free water for a total volume of 15 μL per reaction. **Note:** The qPCR is run in a 384-well plate. However, the reaction volume can be scaled-up for a 96-well plate.
  b. Run a normal qPCR program (can pause overnight and store the DNA at 4°C). **Note:** The Fitpoint (**X**) gives you the number of cycles to use for the PCR (usually between 5 to 15). Upon setting the Fitpoint, perform the PCR amplification with UDI primer pairs.
  a. Use 5 μL adapter-ligated DNA (half of your available DNA) + 1 μL xGen UDI Primer Pair (20 μM) + 15 μL Phusion (ThermoFisher) or Q5 (NEB) + 9 μL RNase/DNase-free water.
  b. PCR program: 98°C 30 sec (1 cycle); ***98°C 10 sec***, ***60°C 30 sec***, ***72°C 30 sec (X cycles determined previously by qPCR);*** 72°C 5 min (1 cycle) 4°C infinite
  **Note:** Do the same for Input samples, except: do X-1 cycles, then add another 1 μL of the same xGen UDI Primer Pair and run one additional cycle of amplification (in order to avoid double peak on bioanalyser). **Note:** For each sample, record the UDI ID for the adapters used as this is required for demultiplexing and assigning samples. **Note:** For NovaSeq and other patterned flow-cell instruments, unique dual indexes are recommended to minimise index hopping and sample misassignment.
36. Post-PCR two-sided library size selection. To each 30 μL of amplified DNA:
  a. Add 70 μL of EB buffer.
  b. Add 65 μL of Ampure beads.
  c. Vortex, spin down quickly and incubate for 15 min at room temperature.
  d. Apply to the magnet, wait for 2 min or until the liquid clears completely and take <u>*the supernatant*</u> into a fresh tube.
  e. Do a 1:1 ratio in the remaining supernatant.
  f. Add 100 μL EB (200 μL total: 30 +70 +100).
  g. Add 135 μL of beads slurry, vortex, spin down quickly and incubate for 15 min at room temperature.
  h. Apply to the magnet, wait for 2 min or until the liquid clears completely and <u>*discard the supernatant*</u>.
  i. Wash two times with fresh 80% EtOH, and dry for ∼10 min (beads should start “cracking”).
  j. Add 11 μL EB buffer, vortex and incubate for 5 min at room temperature.
  k. Apply to the magnet, wait for 2 min or until the liquid clears completely and <u>*take the supernatant*</u>.
  l. Quality control of the libraries are carried out on a Agilent Technologies BioAnalyzer or Tapestation (see expected results in **Figure 3**).
  m. Sequence the libraries using paired-end sequencing with a read length of 150 bp on an Illumina-compatible platform (e.g. NovaSeq, NextSeq).
  **Note**: Always use unique dual indexes (UDI) to mitigate index hopping. **Note:** Configure the NovaSeq run as duall1⍰indexed and use IDT’s UDI index sequences (i7 and i5) for each well in the sample sheet.
37. Running the DRIP-seq bioinformatics analysis pipeline:
  a. Prepare the input files. Populate the `samplesheet.csv` file with all relevant sample information, including: `sample_id, fastq1, fastq2, species, treatment, replicate, tissue, and condition`. Use absolute paths. Make sure that paths are accessible, and file names are correct.
  b. Configure pipeline parameters. Complete the required fields in the `pipeline_parameters.yml` file, including the correct adapter sequences. Double-check that all directory paths, reference genome locations, and other parameters accurately reflect your analysis setup. **Note:** The user can replace the adapter sequence based on the used kit.
  c. Set up the computing environment. The pipeline is written in Nextflow and can run either on a personal workstation or your institute’s computing cluster. Before running, make sure Nextflow and all software dependencies are properly installed (follow GitHub instructions to achieve this). **Note:** If using a cluster, ensure that the architecture and executor configuration in the `nextflow.config` file match your cluster’s current setup. You only need to configure your cluster once. Nextflow will reuse these settings for future runs. Consult your IT team if unsure.
  d. Run the pipeline. Execute the pipeline following the step-by-step instructions on the project’s GitHub page: https://github.com/MCrossleyLab/DRIP-seq_STAR_Protocol
  e. Monitor progress. Nextflow automatically logs pipeline progress and organizes intermediate files. If errors occur, refer to the log files or the troubleshooting section below.
  f. Locate the output files. Output files will be saved in the directory specified under `outdir_path/outdir_name` provided in the `pipeline_parameters.yml` file. This folder will contain subdirectories for different types of results.
38. Exploring output files:
  a. View normalized signal tracks. Open the folder named `normalized_BigWig_files` within your output directory. These files show CPM (Counts-Per-Million) normalized read coverage across the genome.
  b. Visualize results in IGV. Use the free Integrative Genomics Viewer (IGV) to explore the BigWig files. IGV allows you to compare tracks, inspect coverage, and visualize signal patterns.
  Note: Ensure IGV is installed and running on your machine before opening files. Download and installation instructions can be found at https://igv.org/.
39. Downstream analysis:
  a. Access additional output files. All BAM files and other pipeline outputs can be found in the same `outdir_path/outdir_name` directory.
  b. Perform further analyses. Follow the published guidelines in Puzzo F, Crossley MP et al^1^ for recommended downstream analysis steps and interpretation of DRIP-seq results.

**Figure 2:**
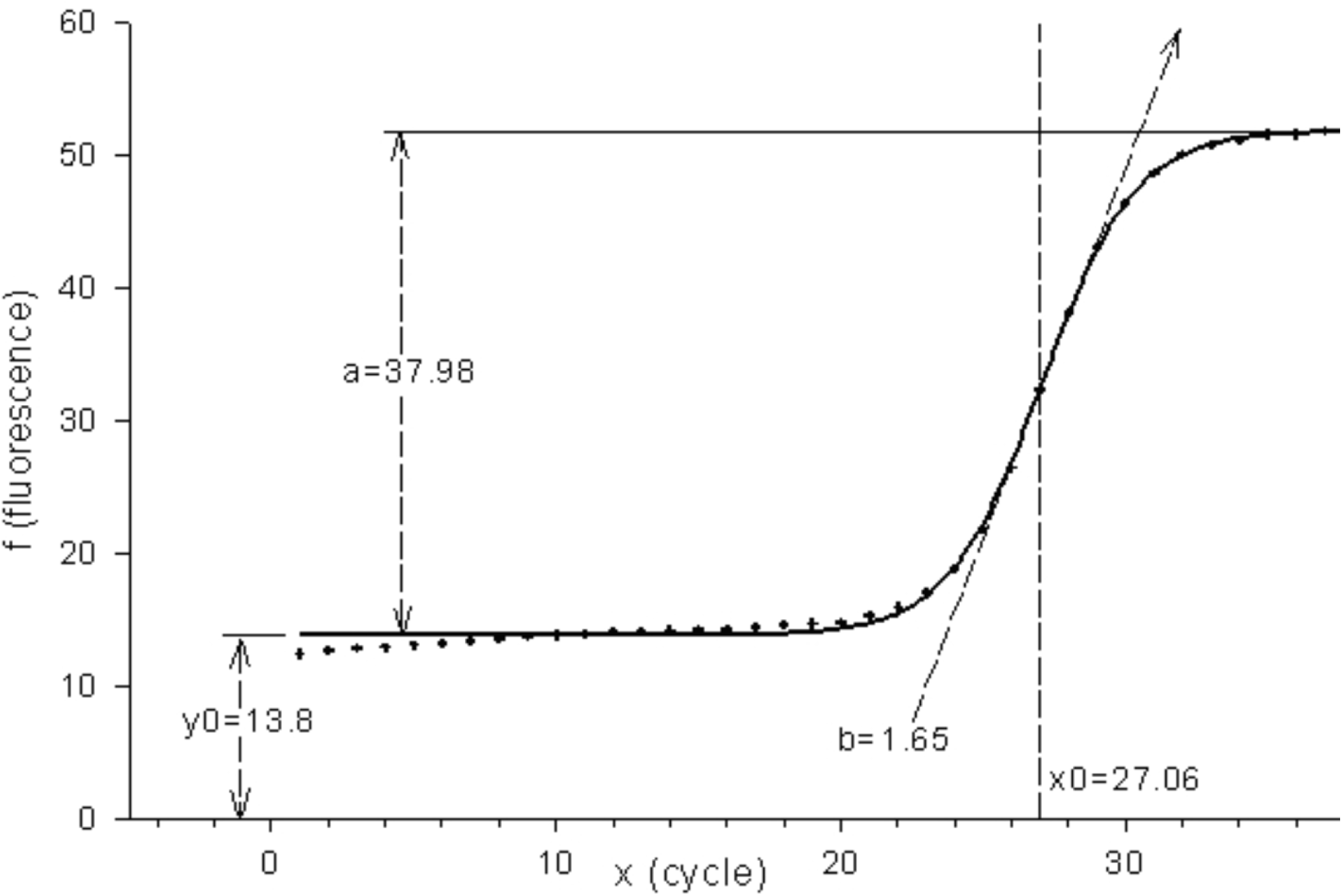
Library preparation overview and calculation of fitting point for library amplification. qPCR curve analysis to calculate the number of cycles for the library amplification by PCR.

**Figure 3:**
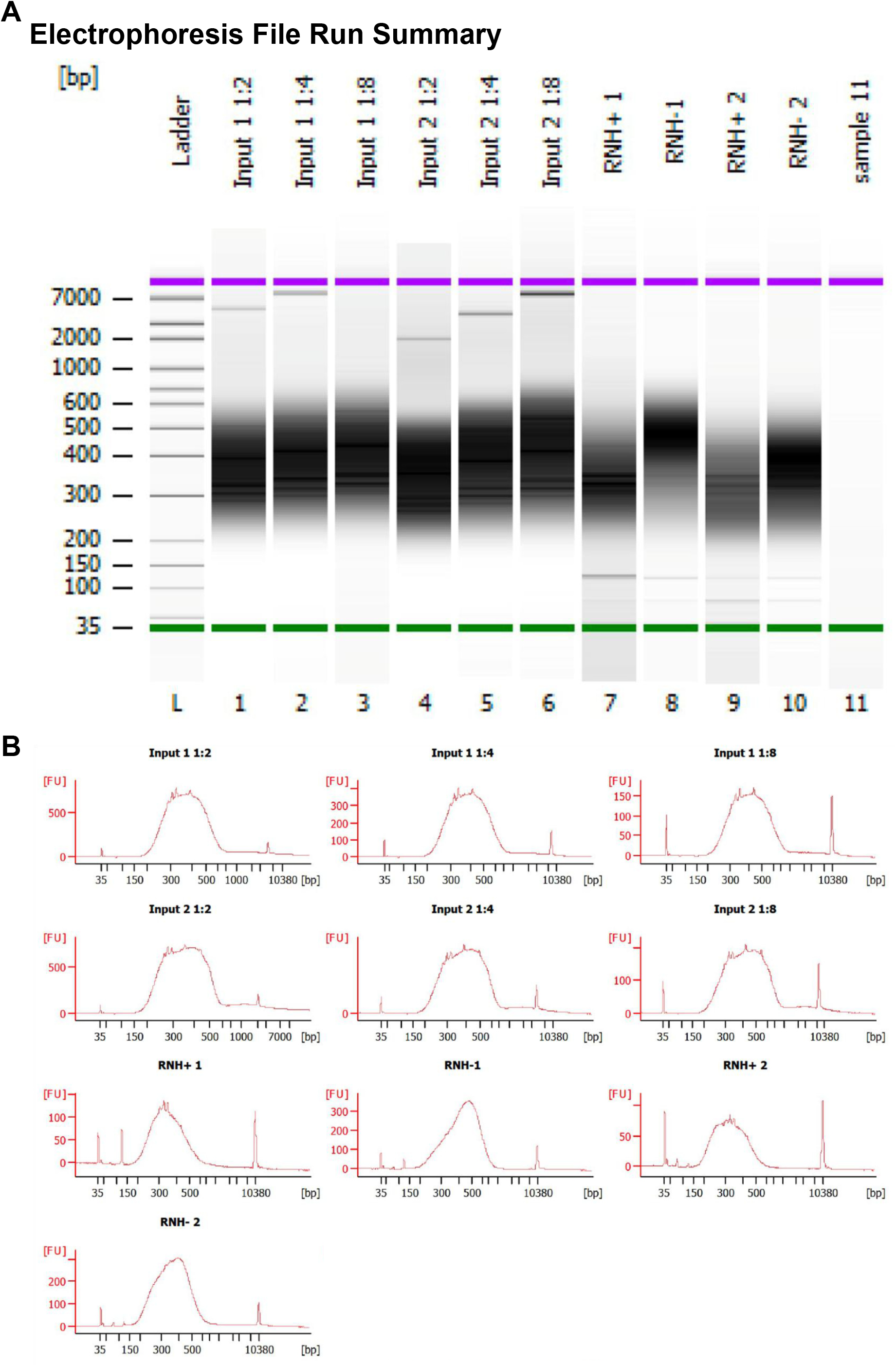
Library quality control. Analysis of the PCR-amplified libraries by Bioanalyzer.

### Expected outcomes

After tissue lysis, genomic DNA is isolated via phenol:chloroform extraction and can be collected in the aqueous phase of homemade phase lock tubes, as described in Step 6. Successful phase separation is shown in **Figure 4**. Samples are fragmented by digestion with a cocktail of restriction enzymes, and DNA-RNA hybrids are immunoprecipitated using the hybrid-specific S9.6 monoclonal antibody. Our protocol results in libraries with a peak of 200-500bp in fragment length, which are ready for DRIP sequencing (**Figure 3**).

**Figure 4:**
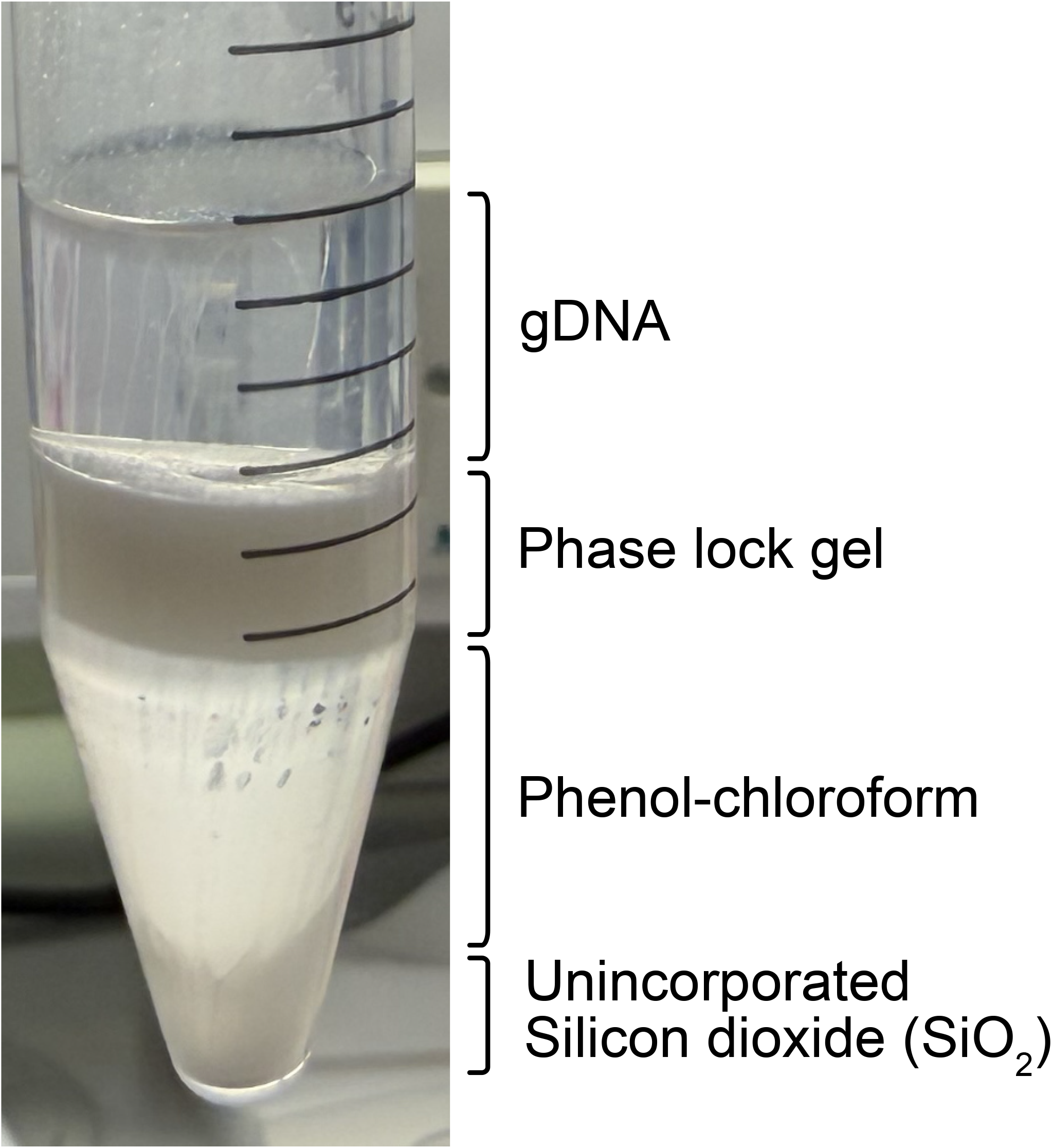
Expected phase lock tube separation of genomic DNA. Phase separation in 15 mL phase lock tubes (following step 6). Aqueous phase containing genomic DNA separates above the phase separation gel.

A successful run of the analysis pipeline should take ∼1h (using the configurations in the `nextflow.config` file).

Following successful sequencing of libraries and alignment to the mouse genome, browser tracks of DRIP-seq signal can be viewed (**Figure 5**). Expect DRIP-seq signal to be robust over genes, including promoters and gene terminators, especially those that are highly expressed in liver (including *Sox9, Alb, Ftl1, Hamp*) (**Figure 5**). IP samples should show enrichment above input samples. Samples pre-treated with RNaseH to remove RNA-DNA hybrids are expected to show reduced DRIP-seq signal genome-wide.

**Figure 5:**
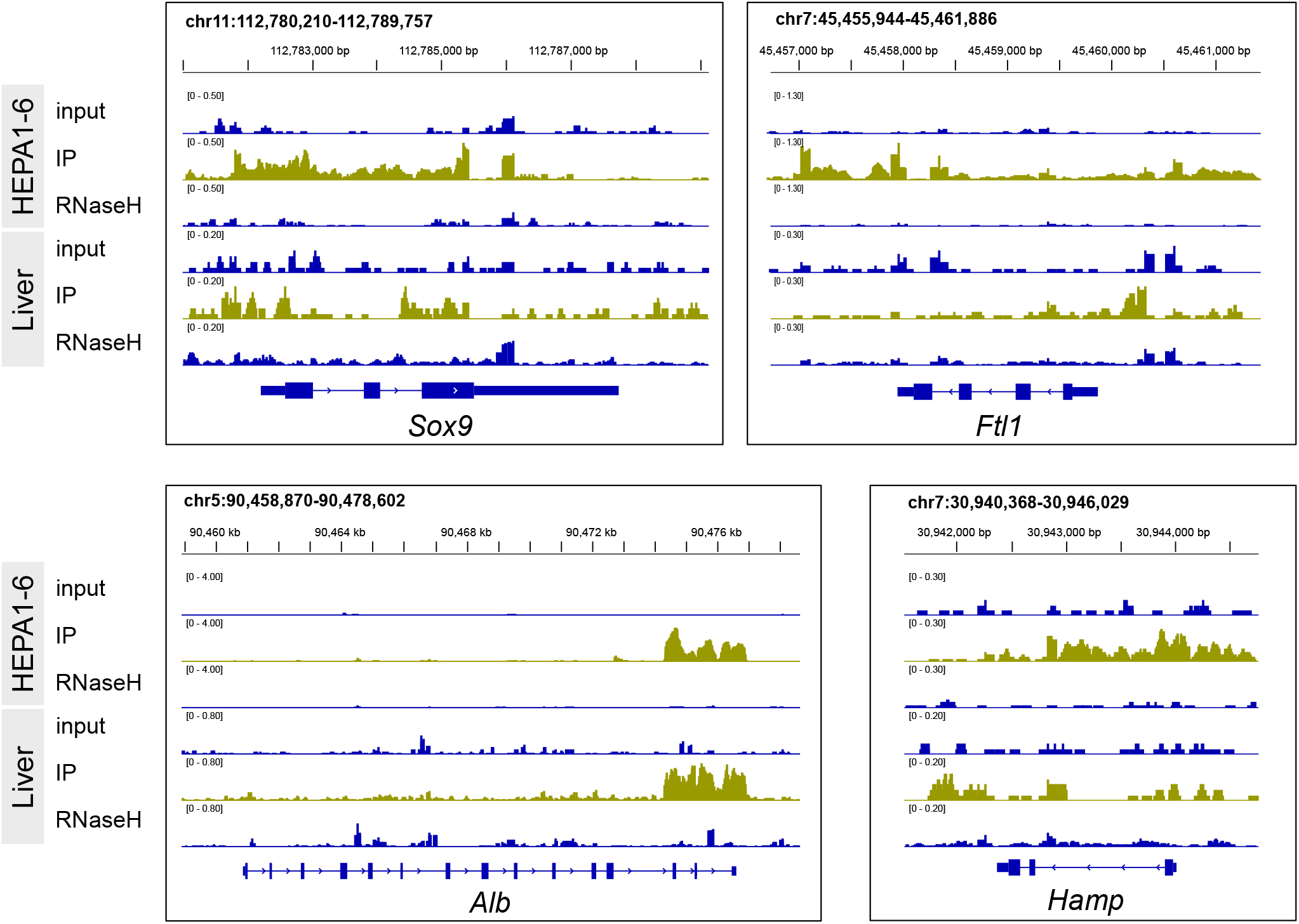
Genome browser tracks of normalized BigWig files from the bioinformatic pipeline. BigWig tracks for 4 representative genes (*Sox9, Ftl1, Alb, Hamp*) are shown as IGV browser snapshots. Tissue identities and sample treatments (input, IP, RNaseH) are indicated on the left. IP tracks are highlighted in green.

### Limitations

One limitation of this protocol is that snap-frozen tissue is processed in bulk. In the liver, DRIP profiles were comparable whether hepatocytes were isolated from the whole organ prior to genomic DNA extraction or the tissue was processed directly after snap-freezing^1^, consistent with the predominantly hepatocytic composition of this organ. Although whole brain tissue can also yield robust DRIP signal^1^, processing bulk material precludes resolution of cell type-specific R-loop landscapes (for example, astrocytes or neurons versus oligodendrocytes). This protocol is therefore most appropriate for DRIP-seq at the whole-tissue level rather than for defining DNA-RNA hybrid formation in discrete cell populations within heterogeneous tissues. While we have validated the procedure using frozen mouse liver and brain, additional optimization and adaptation will likely be required to enable R-loop mapping in other mammalian tissues.

### Troubleshooting

#### Problem 1

##### Incomplete liver homogenization

Liver tissue homogenization may be inconsistent depending on the homogenization system employed. Variability has been observed across instruments, particularly when using bead-based systems (e.g., Next Advance homogenizer with RINO tubes containing metal beads). In such cases, differences in tissue composition among samples, such as varying degrees of fibrosis or lipid accumulation, can affect homogenization efficiency. Fibrotic or lipid-rich livers may exhibit increased resistance to mechanical disruption, resulting in incomplete homogenization.

#### Potential solution

To improve homogenization efficiency, samples may be subjected to additional homogenization cycles until no visible tissue fragments remain. Alternatively, increasing the homogenization intensity (e.g., speed or duration) may enhance tissue disruption. Pre-processing steps may also improve outcomes; liver tissue can be subdivided into smaller fragments at the time of collection or prior to snap-freezing, thereby facilitating more uniform and efficient homogenization during subsequent processing.

#### Problem 2

##### DNA fragmentation following centrifugation of the homogenate

Significant DNA fragmentation may be observed following centrifugation of the tissue homogenate and subsequent resuspension in TE buffer. One contributing factor is insufficient washing of the homogenate with DPBS, which may result in the persistence of cellular debris, including lysed cells and subcellular components. This is typically indicated by a cloudy supernatant after centrifugation. The presence of such contaminants can adversely affect DNA integrity and compromise downstream extraction and analysis.

#### Potential solution

To minimize DNA fragmentation and improve sample quality, the homogenate should be thoroughly washed with DPBS. If the supernatant remains cloudy after the two wash steps, additional washing cycles are recommended. During each wash, the supernatant should be carefully aspirated and replaced with fresh DPBS until it becomes visually clear and a well-defined pellet is observed at the bottom of the tube. This ensures effective removal of residual debris prior to DNA extraction.

#### Problem 3

##### Low yield after immunoprecipitation due to ethanol contamination (related to Step 9)

If pellets are insufficiently dried, contaminating ethanol can reduce restriction digestion and immunoprecipitation efficiency.

#### Potential solution

Remove remaining ethanol with a P10 tip. Allow the pellet to air dry and only proceed when all ethanol drops have disappeared and the pellet is transparent.

#### Problem 4

##### Insufficient genomic DNA recovered from spooling (related to Step 7)

Genomic DNA may be incompletely precipitated or lost due to mechanical shearing.

#### Potential solution

Invert the tube thoroughly to ensure all genomic DNA is precipitated. Use gentle inversion to avoid breaking up the DNA into fragments, which can otherwise be lost in subsequent wash steps (Step 8).

#### Problem 5

##### Incomplete phase separation in homemade phase lock tubes (related to Step 6)

Heavy density phase lock tubes are approximated by supplementing the phase separating gel with silicon dioxide. If they are not made properly, the aqueous phase will migrate below the phase lock.

#### Potential solution

The gel is supplemented with at least 10% silicon dioxide (SiO_2_), but due to variance in particle size, this may be insufficient for good phase separation. Increasing to 15% SiO_2_ may help offset this difference. Alternatively, silicon dioxide can be size fractionated before adding it to the gel. To do so, suspend 10 g SiO_2_ in RNase/DNase-free water and let the larger particles settle for 2 h. Transfer the supernatant to a new tube, then centrifuge to collect remaining SiO_2_. Dry before use.

#### Problem 6

#### Potential solution

Verify that all steps outlined in the GitHub repository have been followed carefully. We highly recommend reviewing the error messages to identify the specific issue and guide troubleshooting. Ensure that the Nextflow run command is executed from the correct environment (see instructions in GitHub). Check that all file paths specified in the `pipeline_parameters.yml` file are accurate and absolute.

#### Problem 7

#### Potential solution

The DRIP-seq pipeline is built in Nextflow, which is designed to parallelize tasks and improve computational efficiency. To achieve optimal performance, it is strongly recommended to run the analysis on a high performance computing cluster rather than a local machine. Before doing so, check that the cluster settings in the `nextflow.config` file are correctly configured and match your cluster’s current architecture. This setup only needs to be done once, and the pipeline will automatically apply the configuration in future runs. For additional guidance, you can refer to the official Nextflow documentation at https://www.nextflow.io/docs/latest/executor.html or consult your IT team for support.

## Resource availability

### Lead contact

Further information and requests for resources and reagents should be directed to and will be fulfilled by the lead contact, Magdalena Crossley.

### Technical contact

Technical questions on executing this protocol should be directed to and will be answered by the technical contacts, Francesco Puzzo (experimental questions -) and Hassan Massalha (bioinformatics questions -).

### Materials availability

This study did not generate new, unique reagents.

### Data and code availability

Original DRIP-seq data files reported in this STAR Protocol are available in Puzzo F, Crossley MP et al.^1^ under GEO accession number GSE261759. Additional code and data analysis files are provided at GitHub: https://github.com/MCrossleyLab/DRIP-seq_STAR_Protocol

## Acknowledgments

We thank members of the Crossley lab for helpful discussions. This work was supported by the Cancer Research UK Cambridge Institute Core Award (SEBINT-2024/100003) to M.P.C. We also thank Matthew Eldridge and Ashley Sawle from the Cancer Research UK Cambridge Institute Bioinformatics Core facility for guidance and support in deploying the analysis pipeline on GitHub.

## Author contributions

F.P. and M.P.C. conceptualized the study. F.P. and C.J.C. designed and conducted experiments. F.P. conducted in vivo DRIP sequencing experiments. C.J.C. developed and optimised phase lock gel separation. H.M. developed and implemented the Nextflow pipeline and performed bioinformatic analysis. All authors wrote the manuscript. M.P.C. provided supervision and funding.

## References

1. Puzzo, F., Crossley, M.P., Goswami, A., Zhang, F., Pekrun, K., Garzon, J.L., Cimprich, K.A., and Kay, M.A. (2024). AAV-mediated genome editing is influenced by the formation of R-loops. Mol. Ther. 32, 4256–4271. 10.1016/j.ymthe.2024.09.035.

2. Gage, G.J., Kipke, D.R., and Shain, W. (2012). Whole animal perfusion fixation for rodents. J. Vis. Exp. JoVE, 3564. 10.3791/3564.

3. Wu, J., Cai, Y., Wu, X., Ying, Y., Tai, Y., and He, M. (2021). Transcardiac Perfusion of the Mouse for Brain Tissue Dissection and Fixation. Bio-Protoc. 11, e3988. 10.21769/BioProtoc.3988.

4. Di Tommaso, P., Chatzou, M., Floden, E.W., Barja, P.P., Palumbo, E., and Notredame, C. (2017). Nextflow enables reproducible computational workflows. Nat. Biotechnol. 35, 316–319. 10.1038/nbt.3820.

5. Martin, M. (2011). Cutadapt removes adapter sequences from high-throughput sequencing reads. EMBnet.journal 17, 10. 10.14806/ej.17.1.200.

6. Langmead, B., and Salzberg, S.L. (2012). Fast gapped-read alignment with Bowtie 2. Nat. Methods 9, 357–359. 10.1038/nmeth.1923.

7. Danecek, P., Bonfield, J.K., Liddle, J., Marshall, J., Ohan, V., Pollard, M.O., Whitwham, A., Keane, T., McCarthy, S.A., Davies, R.M., et al. (2021). Twelve years of SAMtools and BCFtools. GigaScience 10, giab008. 10.1093/gigascience/giab008.

8. Quinlan, A.R., and Hall, I.M. (2010). BEDTools: a flexible suite of utilities for comparing genomic features. Bioinformatics 26, 841–842. 10.1093/bioinformatics/btq033.

9. Ramírez, F., Ryan, D.P., Grüning, B., Bhardwaj, V., Kilpert, F., Richter, A.S., Heyne, S., Dündar, F., and Manke, T. (2016). deepTools2: a next generation web server for deep-sequencing data analysis. Nucleic Acids Res. 44, W160–W165. 10.1093/nar/gkw257.

